# An equilibrium constant pH molecular dynamics method for accurate prediction of pH-dependence in protein systems: Theory and application

**DOI:** 10.1101/2020.11.23.394015

**Authors:** Ekaterina Kots, Derek M. Shore, Harel Weinstein

## Abstract

Computational modeling and simulation of biomolecular systems at their functional pH ranges requires an accurate approach to exploring the pH dependence of conformations and interactions. Here we present a new approach – the Equilibrium Constant pH (ECpH) method – to perform conformational sampling of protein systems in the framework of molecular dynamics simulations in an *N, P, T*-thermodynamic ensemble. The performance of ECpH is illustrated for two proteins with experimentally determined conformational responses to pH change: the small globular water-soluble bovine b-lactoglobulin (BBL), and the dimer transmembrane antiporter CLC-ec1 Cl−/H+. We show that with computational speeds comparable to equivalent canonical MD simulations we performed, the ECpH trajectories reproduce accurately the pH-dependent conformational changes observed experimentally in these two protein systems, some of which were not seen in the corresponding canonical MD simulations.

Table of Contents artwork

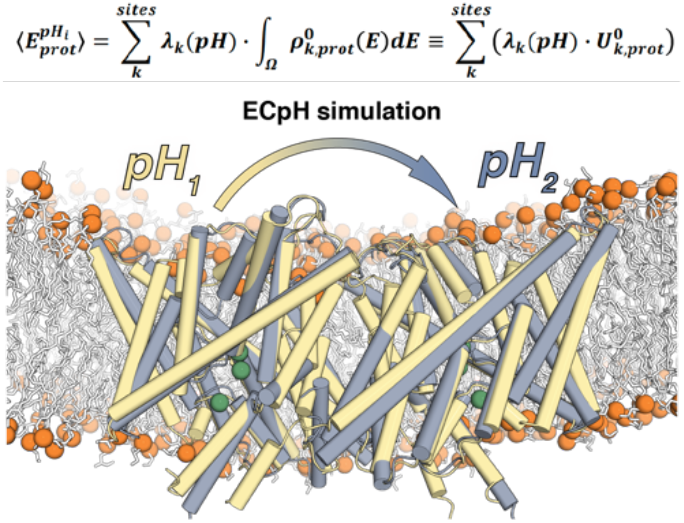

## Introduction

The prospect of exploring the pH dependence of macromolecular biological systems with molecular modeling tools, holds great promise for gaining reliable mechanistic insight and predicting the optimal pH for the peak functional activity. As the functions of a large variety of proteins were shown to be influenced by switching between protonation states of one or more residues^1-5^, an understanding of the atomistic underpinnings of pH dependence should offer powerful tools for regulation of protein activity by increasing the population of specific conformational states, and guiding experimental design.

In canonical molecular dynamics (cMD) simulations the pH of the system is defined by a set of fixed protonation states of all titratable residues that are calculated for a certain pH from per-residue pKa values. However, the pKa values are usually known only for the solvated single amino acids, and not in the structural context of a specific protein where they are likely to be shifted by the non-homogeneous environment. This complicates the assignment of protonation states of protein residues. Prediction methods based on the frozen structure of the protein, such as the used in the popular PROPKA server^6^, have been developed to provide estimates of individual pKa values in a given structure. While this approach is rapid and convenient, it ignores the conformational flexibility of protein macromolecule under realistic conditions, and thereby can miss key elements of the protein’s response to pH changes. The possibility for inaccurate pKa assignments and correspondingly for the incorrect protonation states that it produces for the cMD simulations, can result in inaccurate inferences regarding the states visited by the system and the transitions of interest.

Not surprisingly, the perceived benefits of performing molecular dynamics simulation of proteins with correctly defined protonation states at a given pH have inspired efforts to develop fast and accurate implementations of a number of constant pH (CpH) versions of MD simulations^7-12^. Implicit solvent models have enabled CpH implementation in semi-grand canonical ensemble frameworks, and several newer variations of CpH use allatom simulations that can be coupled to advanced sampling techniques, and sometimes even with polarizable force fields^7,8, 10, 12, 13^.

The approach we present here enables *equilibrium constant pH* (ECpH) simulation within a canonical (or N, P, T) ensemble based on starting with a protein construct in which all titratable sites are fully protonated, and scaling for each site the electrostatic and van-der-Waals interactions using a linear combination of its individual protonation probabilities as the conformation changes in the simulation towards convergence. This procedure takes advantage of a definition of protonation probability for each titratable site and yields a definition of the system in a given conformation at a certain pH. The protonation probability changes accurately with conformation during simulations, as long as the only contribution to the energy difference between two protonation states come from nonbonded interactions. With the model of a given protein state at each pH value obtained in this manner, it is possible to construct the dynamic representation of the pH dependence of a protein from calculated protonation states that reflect the actual proton-occupancies of sites during the dynamics. A major advantage of this development is that it eliminates the approximation of representing protonation states as a binary protonated/non-protonated choice.

We illustrate the new ECpH method and its application for two systems of different size and complexity: the bovine β-lactoglobulin (BBL) known to undergo a pH-dependent conformational change, and the membrane protein CLC-ec1, a Cl−/H+ antiporter in which the dependence on pH leads to major conformational changes involved in regulation of Cl^-^ transport^14-16^. The results are presented in parallel with benchmark tests comparing the ECpH trajectories of these systems to equivalent canonical (cMD) simulations showing that the new method reproduces the stability and ensemble dynamic properties of cMD trajectories. Using direct comparisons with structural data for the two systems to probe the validity of the ECpH results of calculations for a range of pH values, we show that ECpH simulations achieve better agreement with experiment in every detail. For BBL, ECpH reproduces the pH-dependent rigidity of its secondary structure^17-20^, and samples the intermediate conformation observed using tryptophan fluorescence at pH 7^21^, whereas the corresponding cMD simulations miss the intermediate conformations detected experimentally. Similarly, application of ECpH to the larger CLC-ec1 transporter, illustrates the successful representation of the more complex pH dependence of functionally important conformational rearrangements occurring in a pH range of 4.5 to 7.5, and the ability of this new method to bring to light some major repositioning of transmembrane helices inferred from experimental measurements. Thus, recently published experimental data^22^ identified the same changes in the switch from the inward facing conformational state at pH 7.5 to an outward facing opened state at pH 4. Here the enhanced sampling of pH-dependent conformations in CLC-ec1 system is achieved by coupling the ECpH approach to the same Hamiltonian Replica Exchange Molecular Dynamics (H-REMD) protocol that offers promising scalability, accuracy, and significant acceleration of sampling with CpH methods^12^.

## Results

### I. Theoretical framework

Together with the proper assignment of individual pKa values, the suitable representation of the partial nature of protonation state is essential for the accurate prediction of pH dependence of protein structure and dynamics. In the semi-grand canonical ensemble framework, the concept of co-existing protonation states is realized through discrete switching among protonation states. Because of the high frequency of proton exchange, this approach requires significant computational resources, and for the real-world all-atom protein systems remains mostly unfeasible. Here we show that the ECpH method formulated in the same *n,P,T-* ensemble framework commonly used in MD simulations, and using the same information for input pKa values, yields better insight than cMD into the pH-dependence of protein dynamics.

#### The protonation scheme

A central element of the theoretical framework of the ECpH method described here is the construction of the *dynamic protonation scheme* for a molecular system composed of any number of protonatable sites. To enable the shift between protonated and not protonated states, the number of particles is kept the same in the two states, which is achieved by positioning dummy-proton representations at each of the positions that are potentially protonated. The presence/absence of a proton at each site is represented by linearly scaling the non-bonded interactions of sites with titratable hydrogens by an individual *protonation probability* value that is dynamically determined by (1)-the pKa value of that site in a particular state of the entire system, and (2)-by the ambient pH. For a gradual shift between the protonated/non-protonated states of the system, the charges of the atoms neighboring the titratable residues are also scaled linearly, by the same scaling factor. Thus, the core idea of the ECpH method is the introduction of the *modified force field expression* described below, representing the pH-dependence of the state of the system.

#### Expression of the pH dependence of the state

The implementation of this core idea of pH dependence a *canonical ensemble framework* is illustrated first by considering a system with a single protonation site. The protonation probability λ for this case is equal to 1, corresponding to a system at very low pH. Expressing the average energy of the subset of atoms of a titratable residue in this protonated state from the probability density 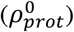 integrated over configuration space Ω we get:

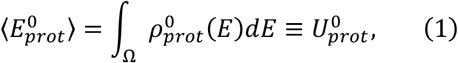

where 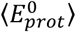 is an average total energy of the single protonation site of the system, 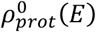 is a Boltzmann probability density of a protonated site over its whole conformational space, and 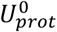 is the internal energy of this part of the system.

The expected pH-dependent differences in dynamics arise from the protonation probability densities modified by the change in proton availability defined by the pH. This pH dependence is introduced through individual protonation probabilities for each protonatable site throughout the system. We assume that the system visits a shared configurational space at the different pH values; degradation processes (e.g., complete unfolding or fragmentation) are outside the scope of this method. Therefore, for a system in any protonation state *i* with a probability density 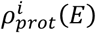, the average energy is expressed as:

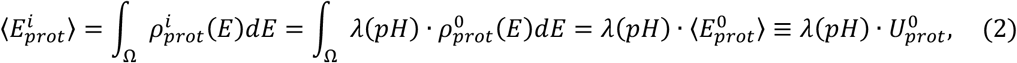

where the probability density function is defined on the same configuration space Ω. Thus, by scaling the internal energy 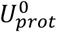 by the λ corresponding to the pH, eq. 2 defines the Boltzmann probability density for any singlesite system with protonation probability λ at a given pH, in terms of the probability density of the fully protonated system 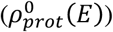.

This definition is equivalent to splitting the entire ensemble into a non-protonated 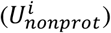 and a protonated fractions 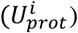:

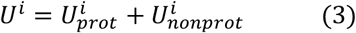

so that at extreme λ values the internal energy would be:

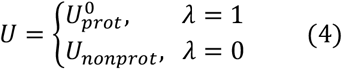

and for any other λ the internal energy of the corresponding ensemble is expressed as:

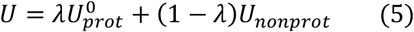

As the protonation probabilities vary at different pH values, the difference between energies of ensembles at different pH values arises from interactions of the titratable proton, so that the internal energies 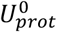 and *U_nonprot_* represent interactions between all atoms of the system and the titratable proton, according to a probability l; this makes *U_nonprot_* equal to zero.

By the same definitions, expressing the average total energy of a system with K titratable sites at a very low pH (*λ*_0_ ··· *λ_k_* = 1) yields:

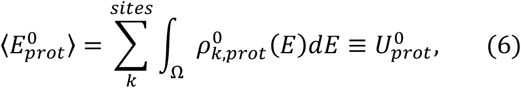

so that for a system with multiple pH-dependent protonation sites and a set of protonation probabilities **λ**(pH):

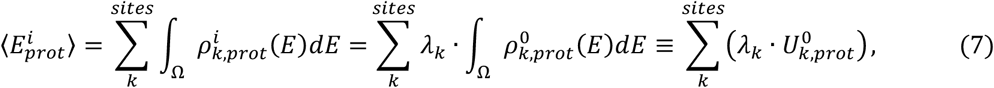

where *λ_k_* is an individual protonation probability for the site k calculated for the current pH. It should be noted that protonation probability values *(λ_k_)* in each state are determined individually for each protonation site by the pH and the pKa. For complex systems with multiple protonation sites, the individual pKa values depend not only on thermodynamical parameters (T, P, n, the nature of the solution), but also on the impact of the other titratable residues and the overall conformation of the system, which can cause shifts of 2-3 units on the pKa scale^22, 23^.

#### Calculations in a given pH range

In the protonation scheme defined by the conceptual framework developed here, a single canonical ensemble is created for each pH in a given range, by using the corresponding protonation probabilities to scale the internal energies of the titratable protonation sites of the system. We follow the common protocol^24, 25^ of applying scaling only to nonbonded terms of the potential energy to eliminate the dependence of kinetic energy on pH. This is achieved by preserving the proton mass in the dummy atoms when λ=0. While this could have a small effect on the dynamics of the system, it does not affect the main goal of proper sampling of the conformation distributions.

#### Definition of the effective forcefield

The electrostatic potential energy and van der Waals terms for interactions of titratable protons are scaled in accordance to the energy definitions in Eqn-s (8) and (9):

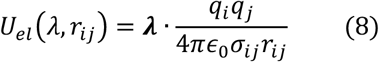

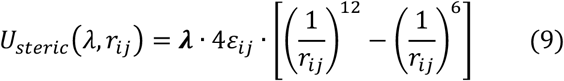

The partial charges of atoms neighboring the titratable residues, which are affected by the change in protonation state are calculated as following:

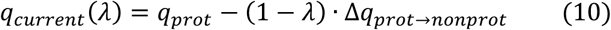

Amino acid residues considered as titratable sites include aspartate, glutamate, histidine, cysteine, and lysine^8^. Because histidine can be involved in two pH-dependent transitions: deprotonation of the protonated state, and transition between the e and δ tautomeric forms, we use two scaling parameters in Eqn-s (8), (9), and (10): one λ to model deprotonation, and a second one – binary – to encode the information about the tautomeric form.

#### Conservation of the net charge

All CpH approaches based on discrete protonation states face a common problem, emerging from the charge-exchange procedure that affects the net charge of the molecular system, which becomes variable, non-zero, during the calculation. Thus, approaches proposed to enable simulations in explicit solvent with PME treatment of long-range electrostatics, were shown to affect the dynamical properties of the system^26, 27^. Here we use an approach that overcomes the problem by distributing the excessive net charge to a sufficiently large number of water molecules in the solution to avoid affecting their state. This is achieved by assigning to the oxygen atoms of randomly selected water molecules an additional partial charge of 0.001e. The extremely dilute acidic environment of this water solution acts as a charge buffer that can absorb from, or provide to, the molecular system the necessary charges without affecting enthalpy, entropy, or the interactions with proteins or membranes.

#### Calculation of the protonation probabilities

Once individual pKa values are set for each protonatable residue in the system, the protonation probabilities are evaluated from the equilibrium constant of protonation/deprotonation process:

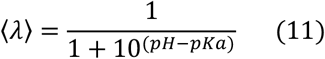

In this paper only several pKa values were set specifically in each system for the key residues that define the pH dependence based on the experimental findings (see Methods section). Otherwise the model pKa-s of singular amino acids in water solution were used.

### II. Performance of the new Equilibrium Constant pH molecular dynamics method

The implementation of the ECpH method to the study of conformational transitions in protein systems is designed to have the general characteristics of conventional MD simulations. This includes the stability of the trajectories generated in the runs, the integrity of the overall protein structure throughout the simulation, and the similarity of dynamic properties of the generated ensembles. To test these elements of the performance of the ECpH method in comparison to parallel canonical cMD simulations, we used as first benchmark a small (162 amino acids) globular protein – bovine β-lactoglobulin (BBL). BBL is known for a signature, pH-dependent conformational shift of the EF loop at pH ~7.3 (Tanford transition) that is attributed to an unusually high pKa value of Glu89 ^4,28^. In addition, the initiation of an unfolding process of BBL at low (< 3) and at high (> 9) pH values, is associated with a series of conformational transitions that have been captured with numerous experimental techniques including NMR, X-ray crystallography and Heteronuclear sequential quantum correlation spectroscopy (HSQC)^21^. We therefore investigated the performance of ECpH method in describing the pH-dependent dynamic properties of BBL from a set of ECpH trajectories obtained at pH conditions ranging from 2 to 11, and compared it to cMD simulations with protonation states of titratable residues varied to represent states at pH 2, 6, and 8.

#### Stability and dynamical properties of ECpH trajectories

The root-mean-square deviation (RMSD) computed along the trajectory for backbone atoms of the protein (see Fig. 1A) is commonly used to infer on the stability of MD methods^12, 29^. We also use the comparison of standard deviation of the RMSD (STD_RMSD_) calculated for short segments of the trajectories to quantify that stability (Fig. 1B). As shown in Fig 1A, the canonical MD and ECpH trajectories initiated from the same conformation of BBL (PDB ID 3BLG, pH ~6.3)^4^ demonstrate common trends of pH dependence. Ensembles generated by the computations at pH 6 refer to the folded conformation of BBL with EF-loop closed and thus should be the closest to X-ray structure. Indeed, at pH 6 the RMSD of Cα-atoms converges rapidly in both ECpH and canonical MD simulations within the first 300 ns, to values falling below 2.5Å on average, with STD_RMSD_ remaining ~0.8 Å (Figure 1B). The initiation of an unfolding process involving loosening of the proteins’ loops was observed at pH 2 in NMR and other spectroscopic studies^18, 30, 31^. Indeed, the RMSD profile of the ECpH trajectory at pH 2 indicates a conformational change of the loops AB, BC, CD, and DE into a loose opened conformation at ~600 ns (Figure S1). In the canonical MD simulations, the geometry of BBL evolves gradually to the same RMSD values. Transition to a more basic environment (pH 8) forces the system into a gradual conformational change associated with the Tanford transition of loop EF.

**Figure 1.**
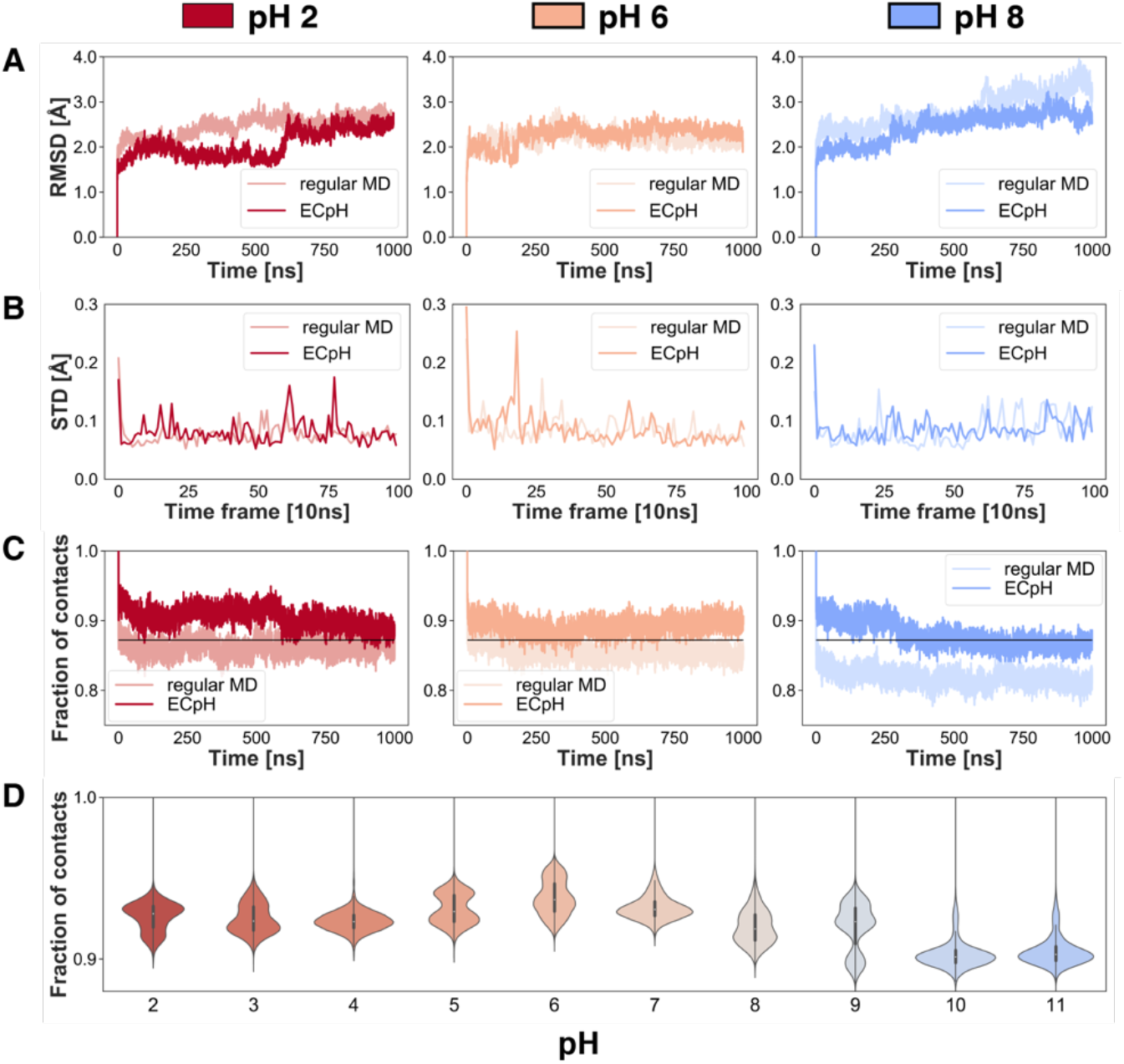
Comparison of results from ECpH and cMD simulations of BBL at pH 2, 6, and 8. (**A**) RMSD of Cd-atoms in 1 μs trajectories at pH 2, 6 and 8 referenced to the X-ray structure at pH 6.3 (PDB ID 3BLG). (**B**) The STD_RMSD_ of these trajectories computed for 10 ns time intervals. **C**) The fraction of native contacts calculated for Cα-atoms with 5Å cutoff. (**D**) The distribution of preserved native contacts from ECpH simulations in the pH range 2-11.

We used a standard technique for testing protein stability, native contact analysis^29^, to evaluate the stability of BBL in the ECpH and cMD simulations, with reference to the crystal structure of BBL at pH 6 (PDB ID 3BLG) in which the EF loop is in the closed conformation (Figure 1C). Interestingly, the changes in the fraction of native contacts in response to pH changes are similar, and both methods reproduce the experimental structural data^17-21, 30-31^ regarding the extremely stable secondary β-barrel structure of the BBL core at the pH range studied which was also confirmed by Solvent Accessible Surface area distributions computed at similar pH range (Figure S2). In more basic environments (pH > 8) a more dramatic loss of native contacts is observed, while the occupancy of the initial state populated at lower pH, vanishes (Figure 1D).

#### Water distribution

Since the force field modification scheme uses explicit water molecules as a charge buffer, the method was also tested for the ability to reproduce proper water dynamics. The radial distribution functional (RDF) analysis demonstrates identical water densities in canonical MD and ECpH trajectories. The water distribution around titratable protons calculated separately, morphs gradually between non-protonated and protonated state densities observed in canonical MD as the pH changes (Figure S3).

#### Computational speed

For practical use of ECpH with real-world systems, the computational speed needs to remain within the range of canonical MD performance. Table 1 shows such a performance comparison for BBL and CLC-ec1 in explicit environments (aqueous, and membrane, respectively), as described in the following section.

**Table 1.** Computational speed of one replica of ECpH vs cMD run on one GPU NVIDIA Tesla V 100s.

| System | Size (atoms) | Computational speed (ns/day) |  |
| --- | --- | --- | --- |
|  |  | ECpH | cMD |
| BBL | 37,504 | 120 | 176 |
| CLC-ec1 | 238,784 | 27 | 53 |

### III. Advantages of the ECpH method in describing the dynamics of the globular protein BBL and the CLC-ec1 chloride transporter over a large pH range

#### BBL

The pH-dependent conformational change in BBL, captured in both the crystal structure and by NMR, is known as a reversible Tanford transition of the EF-loop at pH 7, associated with a highly elevated pKa value of Glu89 (~ 7.3)^4^ “. Below pH 7 the carboxyl of Glu89 is buried in the hydrophobic core of BBL and the gates formed by the EF and AB loops, are closed.

We observed the opening of the EF-loop at pH = 8 in 1 μs trajectories obtained from both methods (Figure 2, right). The process involves gradual reorientation of residues Glu89 and Asn90 which are supposed to switch places during this transition. First, the EF-loop shifts in a direction opposite from the GH-loop; and then the Glu89 carboxyl group reorients out of the hydrophobic pocket, towards the solution, followed by an inward rotation of the Asn90 sidechain to occupy the space left by Glu89. This mechanism is observed at pH 8 in cMD, and at pH 8 and higher in ECpH simulations.

**Figure 2.**
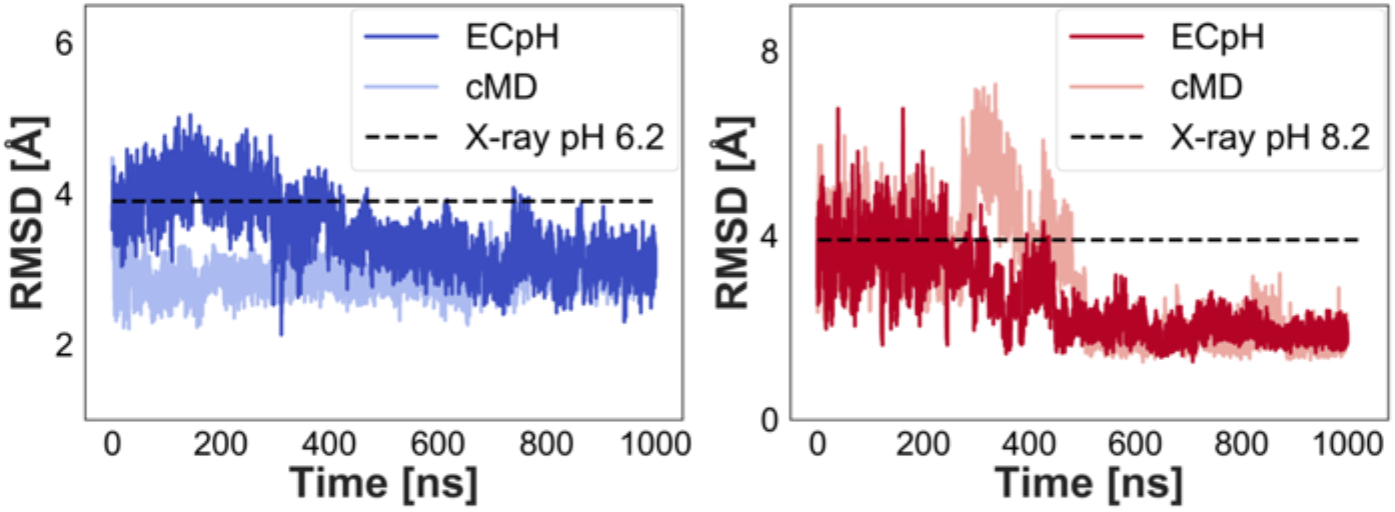
The Tanford transition of EF loop from the closed to opened state at pH 8 (right) and the backwards process at pH 6 (right). The RMSD is calculated for Ca atoms of residues 83-91 as referred to the X-ray structure of the end-point state.

A complete reversal transition into the closed state of BBL at acidic pH was not captured by either ECpH or cMD. Thus, within the 1 μs time frame the backbone of the EF-loop adopts a closed conformation (Figure 2), but the rotation of Glu89 and Asn90 sidechains needed to finalize the transition, does not occur. Notably, however, only the 1 μs ECpH trajectories do sample the beginning of Glu89 reorientation at pH 8, 9 and 10, whereas classical MD simulations remain in the partially closed state throughout.

The distributions of preserved native contacts in ECpH simulations at pH conditions from 2 to 11 (Figure 1D) are consistent with the conformational change occurring at pH 8. Notably, in the ECpH simulation at pH 8 the conformational transition is delayed by ~300 ns compared to the cMD trajectory (Figure 2, left). This delay reflects the fact that in cMD the deprotonation of Glu89 is set artificially, while ECpH represents the more realistic situation of the titratable proton of Glu89 maintaining a ~17% occupancy at pH 8. Notably, the gradual decrease of proton occupancy in Glu89 seen in the in ECpH simulation at pH 7 offers a more detailed description of the mechanism of loop opening, in which the exit of the Glu89 sidechain from the hydrophobic cavity involves first the formation of a hydrogen bond between it and Asn109 (Figure 3). The stable interaction between the EF and GH loops forces the latter to shift and provide space for repositioning of the AB loop (Figure 3A). AB and EF form the gates to the hydrophobic core of BBL.

**Figure 3.**
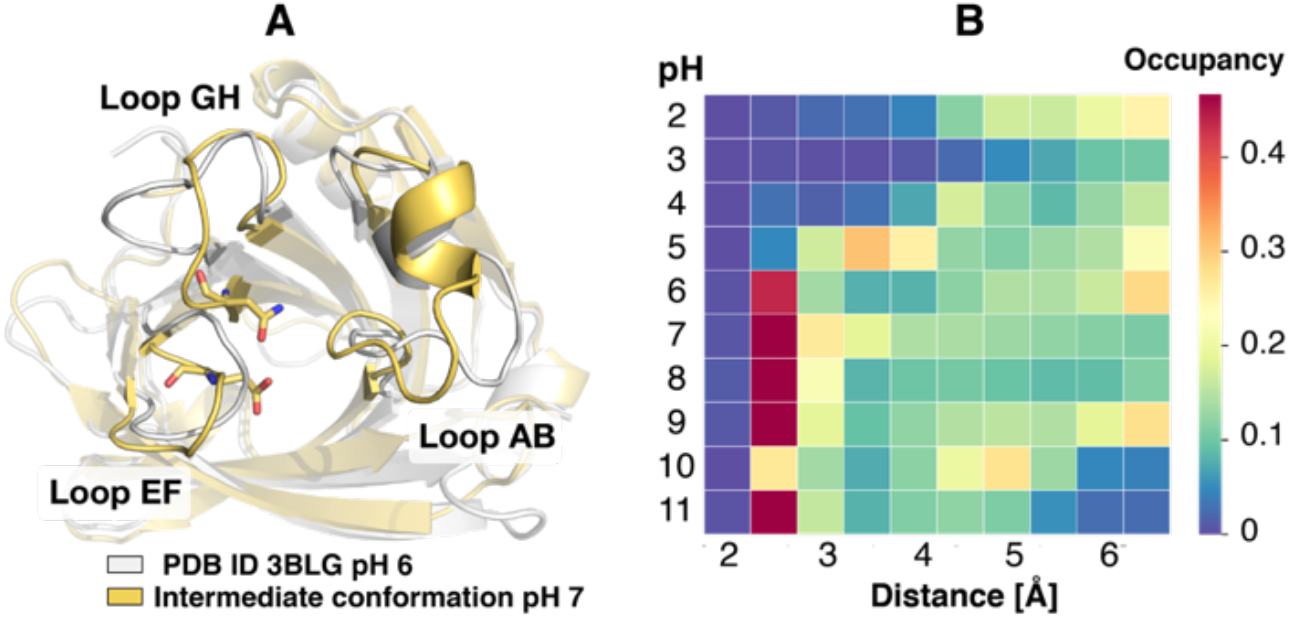
The intermediate state in the BBL unfolding process triggered by Tanford transition at pH 7. **A)** The structure of intermediate conformation is shown yellow, superimposed on the crystal structure at pH 6 (in grey). Glu89 and Asn109 are shown in stick rendering with functional colors. **B)** The distribution of the distance between the carboxyl group of Glu89 and the N atom of the carboxamide group of Asn109 calculated with ECpH in the 2-11 pH range.

The overall conformational change triggers a new orientation of the Trp61 side chain that is now stabilized by π-interactions with Arg40 located on AB (Figure 4A). Notably, the move of the AB loop is observed only in ECpH and not in the cMD simulations. The reason is that in the cMD runs, the dependence of the Trp61 sidechain on pH, is lost. In contrast, the ECpH trajectories show that Trp61 visits three orientational states. State 2 (Figure 4B) is sampled at pH 7 and reflects the conformational changes discussed above. State 3 is visited only very briefly in cMD at pH 6 (Figure 4C), whereas in the ECpH simulations it is strongly populated at extreme pH values at which multiple experiments observed the beginning of unfolding. Thus, in a basic environment (pH > 9.5) the fraction of secondary β-sheet structure of BBL was shown to decrease rapidly while the portion of disordered protein structure increases^19^. Although the unfolding process is too slow to be captured in the 1 μs simulations we carried out^18^, the initiation of the process is revealed in the mechanistic details emerging from the ECpH simulation.

**Figure 4.**
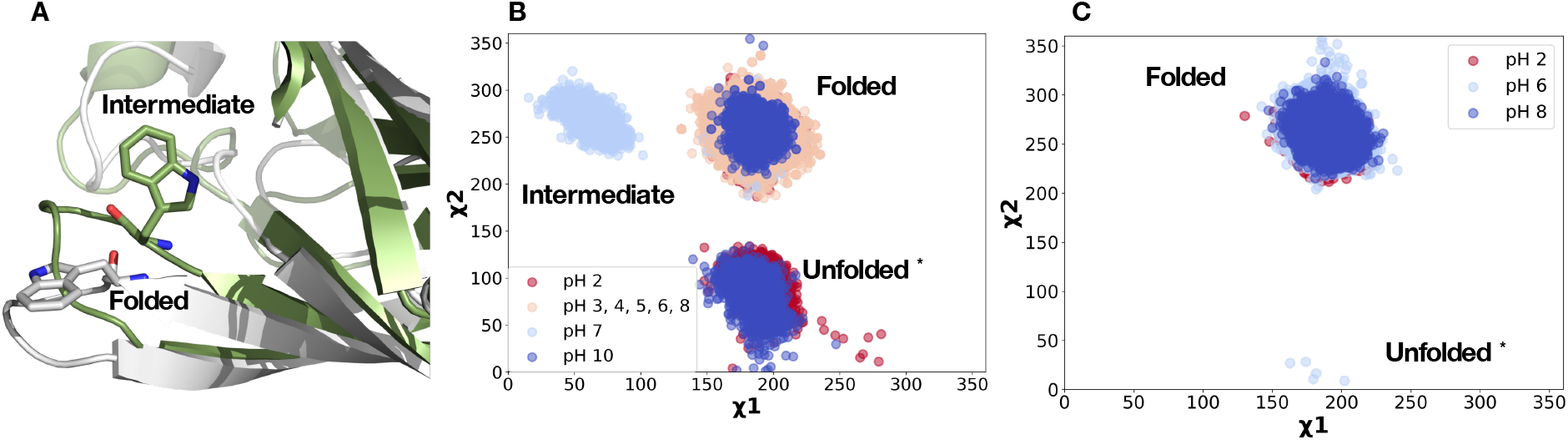
The conformation of Trp61 side chain in three states of BBL unfolding. **A)** Orientation of Trp61 in an intermediate state at pH 7 detected by tryptophan fluorescence. **B)** Distribution of dihedral angles in ECpH at pH from 2 to 11. **C)** Trp61 conformations sampled in regular MD at pH 2, 6, 8.

The three orientational states of the Trp61 (CD-loop) in the ECpH s trajectories are in agreement with experimental results from monitoring the Trp61 (CD-loop) fluorescence at pH ~2 (2.0 – 3.5), ~4 (4.0 – 5.5), ~6 (6.0 – 7.0), 7.5 and 8.0 by Sakurai K. et al.^21^. The three stable states observed were assigned to states of BBL unfolding: native, loose, and intermediate. The intermediate of the unfolding process was attributed to the reorientation of Trp61 which the authors considered to be triggered by the Tanford EF transition occurring at ~ pH 7. The ECpH results in Fig. 4 illuminate the role of Trp61 in the process.

Like the agreement with the major conformational changes observed experimentally, root-mean square fluctuation (RMSF) of the backbone atoms in the cMD and the ECpH simulations (Fig. 5B vs 5C) compare well with NMR data (in Fig 5A) for the BBL protein at pH 2 (PDB ID 1CJ5, 1DV9)^30, 31^. Because the number of NMR ensembles in the PDB records (10 and 21 states, respectively) is limited, the comparison considered only the positions of the peak in the fluctuations of the identified elements of secondary structure. The increased flexibility of loops AB, BC, CD, EF and of the Ha helix observed at pH 2 in the NMR ensembles, is reproduced in the ECpH simulations in acidic environment (pH 2-5). In the cMD trajectories, however, the BBL protein appears to be more rigid than expected at acidic pH, as the fluctuations of BC, CD and Ha either do not differ from those at other pH values or are even decreased.

**Figure 5.**
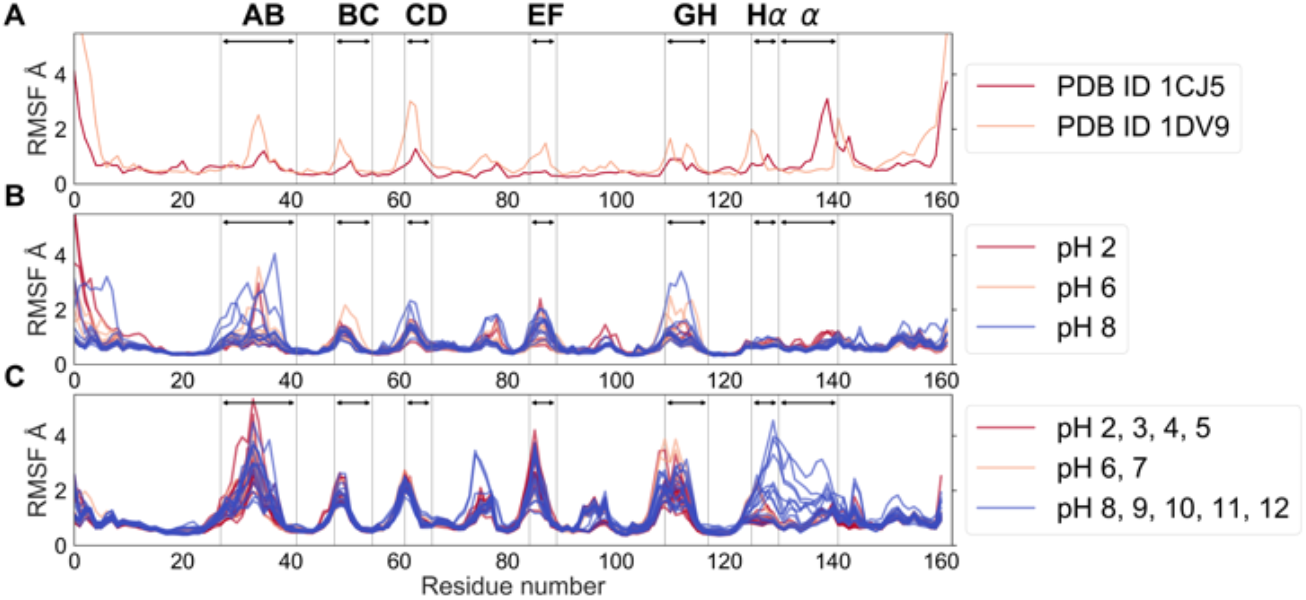
The RMSF of BBL backbone atoms from **A)** 25 NMR states, **B)** regular MD and **C)** ECpH simulations.

Unlike the other structural elements, the increased flexibility of Helix a observed with NMR at acidic pH, is not captured in the either method (Figure 5a), most probably because the trajectories are too short (1 μs) to model the helix unfolding. However, the ECpH method does record the significant mobility of Helix a in the extremely basic environment (pH > 10), in line with the decreased fraction of ordered secondary structure observed at pH > 9^19^. At neutral pH the mobility of all loops is reduces in both MD methods. Notably, the decreased mobility of the AB loop at basic pH compared to acidic and neutral conditions is the consequence of the conformational change detected in the ECpH simulations at pH 7. In cMD, however, the mobility of the AB loop is increased at pH 8, impairing the hydrophobic core integrity that requires the observed closure of the AB loop at that pH.

#### CLC-ec1

The performance of ECpH in the modeling of pH-dependent conformational changes in a class of larger and more complex proteins, in a membrane environment, is illustrated for the CLC-ec1 transporter protein that performs the coupled antiport of Cl^-^ and H^+^^32, 33^. The H+/Cl− exchange follows a ~ 1:2 stoichiometry that suggests a pH-dependence of the transport function^32, 34^. Indeed, the efficacy of CLC-ec1 mediated transport is pH dependent, and the largest Cl^-^ uptake is observed at pH 4.5. Interestingly, crystallography data for the WT CLC-ec1 at pH 4.6 (PDB ID 1KPL) and pH 8.5 (PDB ID 1KPK) does not show major difference in conformational state of the protein^35^. However, other experimental evidence, e.g., from NMR, was suggested to indicate that the interconversion between functional states of the CLC protein in response to pH changes involves large-scale conformational changes such as the movement of Helix R and the structural change of the P-Q linker^36-38^. The introduction of a disulfide crosslink by the mutations A399C/A432C inhibited Cl-transport, leading to the suggestion of a connection between the displacement of Helix O and the gate opening in outward-facing state in WT protein^16^. Major structural rearrangements were also suggested from DEER experiments, cross-linking, and NMR experiments, which concluded that motions of helices G, N, O and P act to widen the extracellular bottleneck^14, 15^. The recent high-resolution X-ray structure of a mutant construct of the CLC-ec1 protein (E148Q/E203Q/E113Q, PDB ID 6V2J) termed QQQ-CLC^14^, does show structural rearrangements to form an outward-facing open conformational state at pH 7.5 that mimics the WT-like conformation at low pH. We therefore used the QQQ mutant structure as a reference to investigate the pH-dependent changes predicted from parallel ECpH and cMD simulations in that pH change.

According to the QQQ-CLC data, the conformational change occurring between pH 4.5 and pH 7.5 involves the repositioning of helices M, N, O, P, and G. This is supported by the inter-subunit distance distribution for Cα-atoms measured in DEER for Met373, Val374 (Helix N), Glu385 (Helix O), Asp417 (Helix P) and Asp171 (Helix G). From the trajectories obtained for the two pH conditions, we evaluated first the distribution of distances between the Cα-atoms of the residues A399 and A432 within a subunit that is believed to change during the chloride transport connected by the presence of a crosslink in the A399C/A432C mutant. The change in the distribution of 399-432 Cα-atoms distances between pH 5 and 8 is evident in the ECpH trajectories, but is absent in the cMD simulations (Figure 6, right). In the DEER experiment the inter-subunit distance of Asp171 which is located on the intracellular side of the CLC protomers exhibits the largest change observed among those detected experimentally (6 Å) and is reproduced by the data presented here from the ECpH calculation, where the second distribution peak of a new conformation appears at pH 5. Meanwhile the distributions obtained with cMD shows only one peak for Asp171 inter-subunit distance at both pH values.

**Figure 6.**
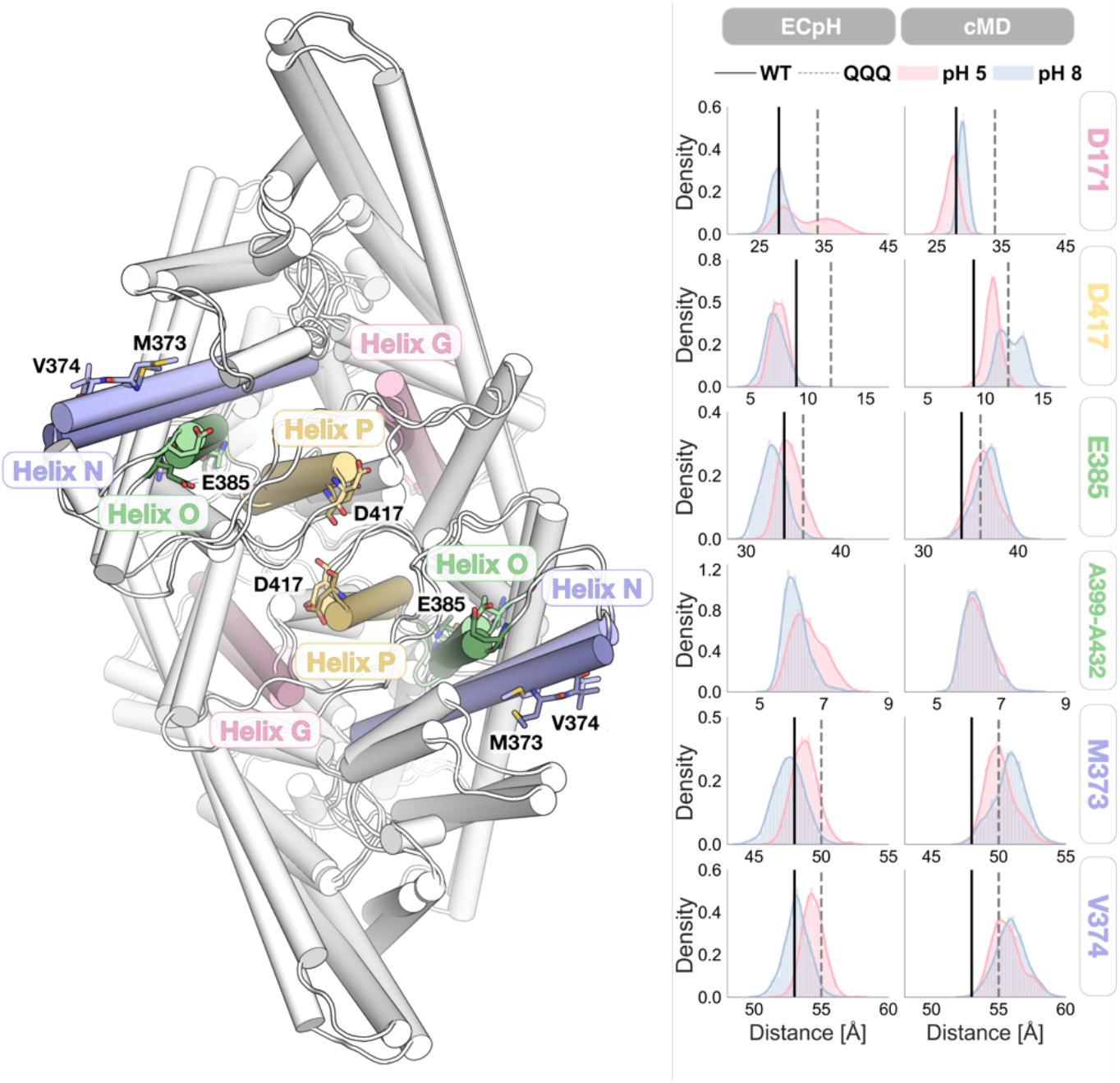
**Left**: the comparison of the inter-subunit distances obtained experimentally with MD data from ECpH, and cMD simulations. **Right**: Detailed representation of the conformational changes undergone by the key structural elements involved in the overall shift.

Other distance changes observed in the DEER experiment are for Met373, Val374, and Glu385, all of which are located at the extracellular end of CLC-ec1 (Figure 6, left). Although subtle, the conformational rearrangement of Helix N recorded in the ECpH ensemble at acidic pH, agrees perfectly with the experiment. Further, the shift in inter-subunit distance distribution for E385 agrees well with the experimental results used to suggest a repositioning of Helix O as suggested in ref. 14. As seen in Figure 6, however, the predictions of pH-dependent relative changes obtained from the comparative cMD simulations at pH 4.5 and 7.5 were not in agreement with the experimental data^14^. Analysis of the likely reason for the discrepancy between the experimental and ECpH-computed results for the Asp417 in the P-Q flexible loop at the CLC-ec1 dimer interface indicates that the larger movement of Asp417 observed in the DEER experiment may depend on a preceding reorientation of Tyr419 (also in the P-Q loop). Indeed, the inter-subunit distance between Cα-atoms of Tyr419 calculated with ECpH (Figure S4) suggests a larger distance between the P-Q loops than that inferred from the Asp417 data. Because it had been shown previously^37-38^ that at pH 4 the orientation of Tyr419 (P-Q loop) changes from buried to outward-facing, it is reasonable to infer from the relative positions of Asp417 and Tyr419 that for the former to move, the latter needs to reorient and clear the space inside the buried area in P-Q region. This movement has started in the ECpH simulations and longer trajectories are likely to eliminate the discrepancy.

The extent of transition to the active state represented by the inter-subunit densities in 0.5 μs H-REMD ECpH simulations at pH 5 (Figure 6), is quantified for each subunit by the distribution of RMSD of G, N, O and P helices (referenced to PDB 1OTS; Figure S5). Of the four helices involved in pH-dependent structural change of CLC-ec1 only helix G shows similar RMSD occupancies for the new state in both subunits, while the displacement of the other helices is not symmetrical in the two. As the CLC-ec1 monomers are known to be functionally independent^35^, the asymmetrical response to a pH change is not surprising, and the same changes are expected to occur in both monomers during longer trajectories.

## Discussion

The new equilibrium constant pH method presented here was designed to enable the exploration of pH dependent structural dynamics of macromolecular systems with greater accuracy and reliability than conventional MD simulations in a framework that conserves their properties and practical attributes. The trajectories generated within ECpH for the illustration of the method and its performance were indeed shown to have the same level of stability as the corresponding cMD trajectories, with similar performance characteristics and dynamics, at similar computational speeds.

The differences in the pH-dependent behavior of the systems described by ECpH simulations compared to cMD, point to the significant advantages of using partial protonation probabilities, over the binary representation of protonation changes in cMD. Because ECpH is able to distinguish in this manner between the effects of pH in a narrow range of physiologically relevant values (5 – 8) without the precise knowledge of all individual pKa values of titratable residues, it informs on the effects of the protein’s dynamics in a range in which only histidine is expected to change its protonation state in the cMD framework. In the BBL system, for example, this shortcoming prevents cMD from detecting the intermediate state observed at pH 7 in the tryptophan fluorescence study. Similarly, in the CLC-ec1 example, extensive cMD sampling at pH 4 does not provide insight into structural changes captured experimentally with DEER, X-ray and NMR, but these are reproduced readily in ECpH simulations.

The binary representation of the protonation states leads to the faster transition observed for the EF-loop in BBL at pH 8 (Figure 1) in the cMD simulations than ECpH. This difference holds until pH 10, where the conformational shift occurs almost simultaneously in both methods (Figure 1D). This reflects the calculation of protonation probability in the ECpH (but not cMD) according to the Hill equation (Eqn. 11). Until pH 10 is reached, the protonation probability of Glu89 is < 0.1. Because the binary representation of protonation states in cMD means that the protonation probability at a titratable site is either 1 or 0, it represents accurately only extreme pH conditions. Notably, in systems with multiple protonation states this may lead to the co-existence of protonation states of neighboring residues that do not actually correspond to the same pH. The accelerated transition of the EF-loop in cMD produced by the binary representation of the protonation state is thus contributing to its inability to reach functionally relevant intermediate states that appear in experiment and the ECpH simulations.

Unlike the direct comparison of the performance our ECpH method with cMD, the comparison with other CpH approaches is challenging because of the paucity of reports of conformational sampling of protein systems with such methods. Still, the CpH simulations are and remain an essential preliminary step in modeling pH dependence of proteins when the pKa values of the residues that define the conformational response to pH change are not known from the experiment. The most recently developed CpH methods enable such calculations with either coarse-grained force fields or full-atom representations using classical or polarized force fields^7-12^. The evaluation of pKa values for all the residues of protein systems of interest remains a computational challenge, however, despite considerable improvements in speed^7, 9^. ECpH lessens this burden somewhat by requiring the knowledge of individual pKa values only for key residues. As shown here in the calculations for BBL and CLC that reproduced a pH-dependent conformational changes in a wide range of pH values, only the pKa values of Glu89 (BBL) and Glu113, Glu148 (CLC-ec1) were specifically set. Because limiting the number of initially set pKa values in the ECpH calculations, the resulting protonation conditions at various pHs are correct only relative to each other in the molecular system, which may skew somewhat the prediction of the absolute pH value at which the transitions occur. For BBL, the pH value at which conformational transitions are observed in the ECpH calculations matched the experimental data in spite of the fact that only the pKa of Glu89 was set. But for CLC-ec1, the processes leading to pH-dependent conformational shifts are more complex and may not be due only to the pKa values of Glu113 and Glu148 which were set. Indeed, both the nature of the conformational transitions in CLC-ec1, and the range of pH values in which they occur in the ECpH simulations, were identified and assigned correctly with respect to the experimental data. However, there was a slight difference between the pH values reported for the conformational from experiments (from pH 7.5 to pH 4.5) and from our ECpH simulations (transitions between pH 5.0 and pH 8.0). While rather insignificant in this particular case, this slight difference indicates a degree of dependence of the results from the ECpH method on the correct choice of the specific residues for which the knowledge of pKa values is necessary.

## Methods

All MD simulations (both ECpH and cMD) were performed with OpenMM 7.3 software^39^ using the CHARMM36 all atom force field^40^, in the NPT thermodynamic ensemble at P = 1 bar and T = 310 K. The pressure was preserved by either a Monte Carlo barostat or its membrane modification implemented in OpenMM (MonteCarloMembraneBarostat)^39^. Long-range electrostatic interactions were evaluated with PME. Nonbonded interactions cutoff and switching distances were set to 12 Å and 10 Å, respectively. The integration step was set to 2 fs while the fluctuations of water bonds were constrained. The list of pKa values used in assigning the protonation states for titratable residues were Glu: 4.4, Asp: 4.0, Lys: 10.4, His 6.5, 9.1, Cys: 9.5, unless stated differently (see below). Cys residues involved in disulfide bridges were not titrated.

**For BBL** the ECpH simulations were conducted for a pH range from 2 to 11, for 1 μs each, and were initiated from both opened and closed states of the EF loop. The crystal structures of BBL at pH ~ 8.1 (PDB ID 2BLG), and at ~ 6.3 (PDB ID 3BLG), were used as starting points. In all the simulations, the pKa value of Glu89 was set to 7.3 according to experimental data^4, 28^.

The corresponding cMD simulations were performed for pH 2, 6 and 8. At pH 2 all titratable residues were protonated, while ensembles at pH 6 and 8 differed in the protonation the protonation states of Glu89 and His residues. The simulations were also performed for 1μs each, starting from both states of the EF loop.

**For CLC-eq1** the simulated molecular construct was built from PDB ID 1OTS^41^ and was embedded in the membrane with the CHARMM-GUI Membrane Builder server^42-43^. The membrane was composed of 629 lipid molecules with a 70:30 composition of POPE:POPG. The system was solvated in explicit water solution with 0.15 M KCl. The equilibration procedure followed the standard CHARMM-GUI equilibration protocol in NAMD 2.10 software^44, 45^.

The ECpH MD simulation was performed for a pH range from 2 to 9. The Hamiltonian replica exchange protocol was applied every 0.1 ns of 0.5 μs ECpH MD simulation to a random pair of replicas. pKa values were set for Glu113 and Glu148 at 8 and 6.5, respectively according to experimental and computational data^5, 46^. All other titratable residues were assigned the pKa values of the corresponding amino acids. For cMD of the CLC-ec1 system data were generated from 2 parallel trajectories of 1.6 μs each for both pH 4.5 and pH 7.5.

### Native contact analysis

The native contact analysis was performed on 1 μs trajectories at a pH range from 2 to 11 for ECpH and pH 2, 6 and 8 for cMD. The distance cutoff was set to 5 Å.

## ASSOCIATED CONTENT

### Supporting Information

The Supporting information contains depiction of a loose loop conformation of BBL protein at pH 2, a Solvent Accessible Surface Area (SASA) distribution for the hydrophobic core of BBL protein, the radial distribution of water molecules around the titratable residues, the RMSD distribution for CLC-ec1 helices G, N, O, P in the two subunits of the dimer and the inter-subunit distances between Tyr419 residues from ECpH simulations at pH 5 and 8. The ECpH python code compatible with OpenMM is published on the GitHub: https://github.com/weinsteinlab/pH-Replica-Exchange

The Supporting Information is available free of charge on the ACS Publications website.

### Funding Sources

The work was supported by National Institutes of Health (NIH) Grant R01GM106717 (H.W.), and NSF award BIGDATA: IA: Collaborative Research: In Situ Data Analytics for Next Generation Molecular Dynamics Workflows (NSF #1740990).

### Notes

The authors declare no competing financial interest.

## ACKNOWLEDGMENT

The authors thank Dr. George Khelashvili and members of the Weinstein lab for helpful discussions. The computational work was performed using the following resources: the Oak Ridge Leadership Computing Facility (INCITE allocations BIP109) at the Oak Ridge National Laboratory, which is a DOE Office of Science User Facility supported under Contract DE-AC0500OR22725; resources services, and support provided at the RPI Artificial Intelligence Multiprocessing Optimized System (AiMOS) system accessed through an award from the COVID-19 HPC Consortium (https://covid19-hpc-consortium.org/), which is a unique private-public effort to bring together government, industry, and academic leaders volunteering free compute time and resources in support of COVID-19 research.

## Notes

https://github.com/weinsteinlab/pH-Replica-Exchange

### Competing Interest Statement

The authors have declared no competing interest.

