## Supplementary material for "An equilibrium constant pH molecular dynamics method for accurate prediction of pH-dependence in protein systems: Theory and application"

### Table of Contents.

S1 Conformation of the AB, BC, CD and DE loops of BBL protein at pH 2

S2 Integrity of the hydrophobic core of BBL protein shown by SASA analysis and by the distribution of Trp19 sidechain conformations at different pH in cMD and ECpH.

S3 The radial distribution functional analysis of water molecules and titratable sites of cMD and ECpH trajectories.

S4 The distribution of inter-subunit Ca distances for Tyr419 of CLC-ec1 transporter at pH 5 and 8 from ECpH simulation.

S5 The distribution of RMSD of G, N, O and P helices from 0.5 $\mu$s HREMD ECpH simulation at pH 5 for two subunits of CLC-ec1 dimer.

Figure S1. Conformation of the AB, BC, CD and DE loops of BBL protein at pH 2

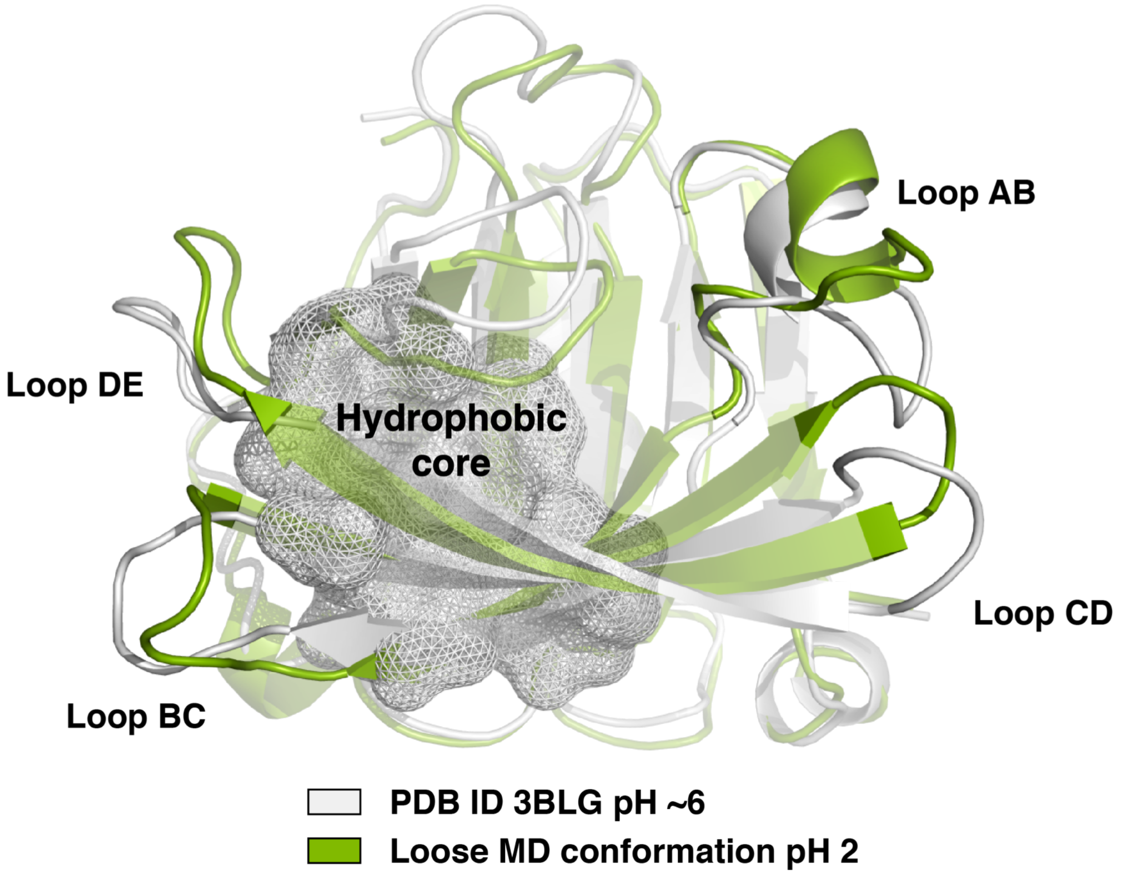

Figure S2. **Upper panel:** Solvent accessible surface area for hydrophobic core of BBL protein at different pH in regular MD simulation (Left) and ECpH MD (Right).The SASA values were calculated in MDtraj python package, with probe radius = 0.14 nm, number of sphere points = 960, in residual mode. **Lower panel:** the distribution of sidechain dihedrals of W19 in cMD (Left) and ECpH (Right) simulations.

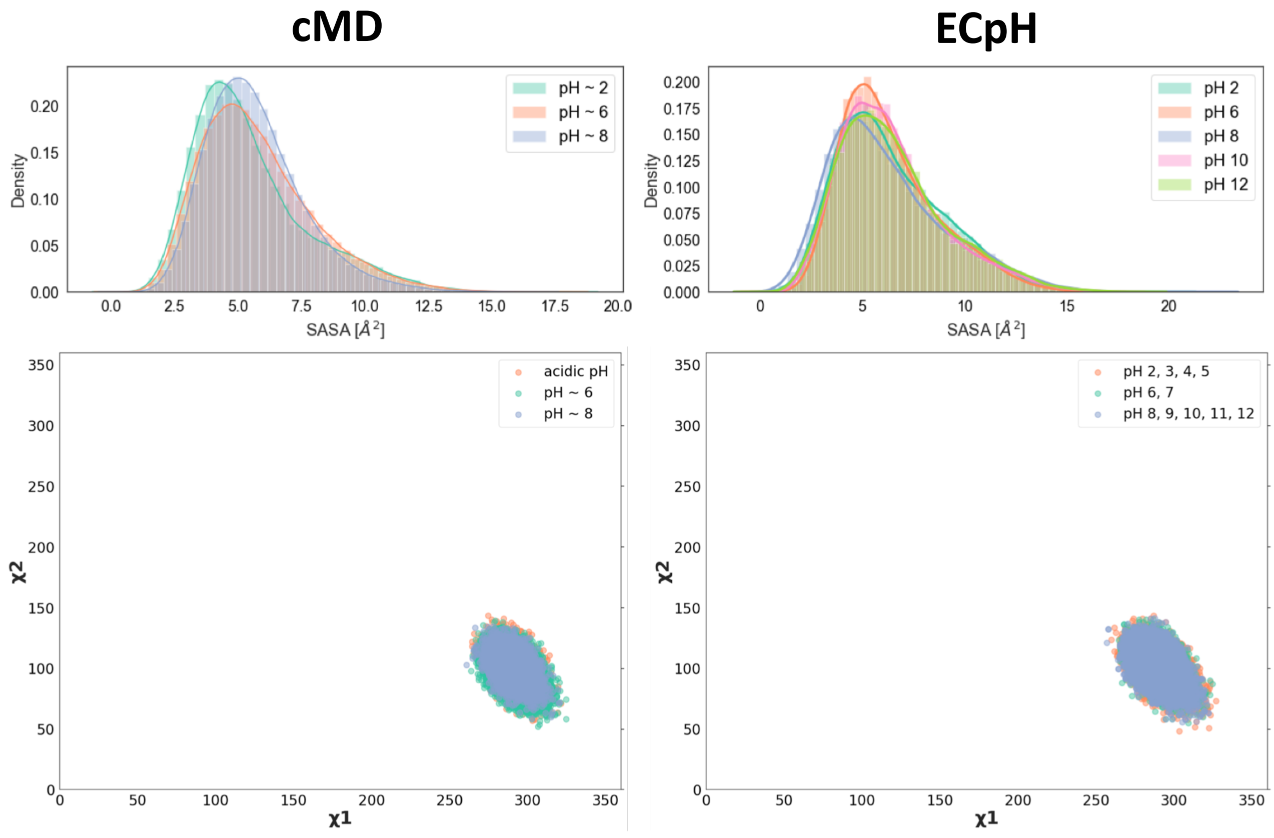

Figure S3. The radial distribution of water molecules, calculated between atoms of the protonation sites and potential hydrogen bond partner in water molecules for ECpH (left) and regular MD (right). The RDF analysis was performed with MDtraj python package, for distance range from 0 to 10 Å under minimum image convention.

**ECpH**

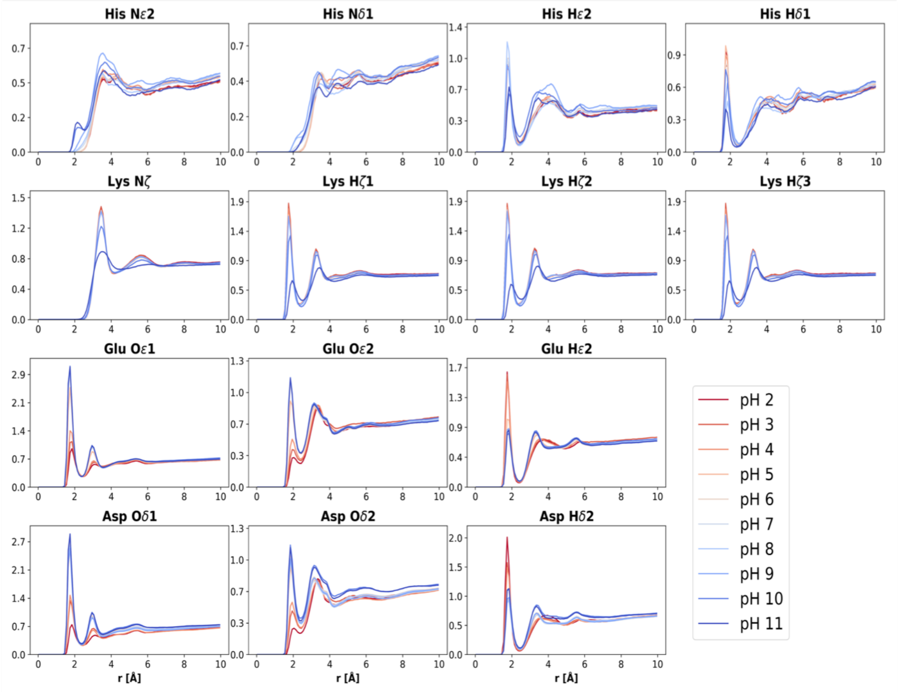


**cMD**

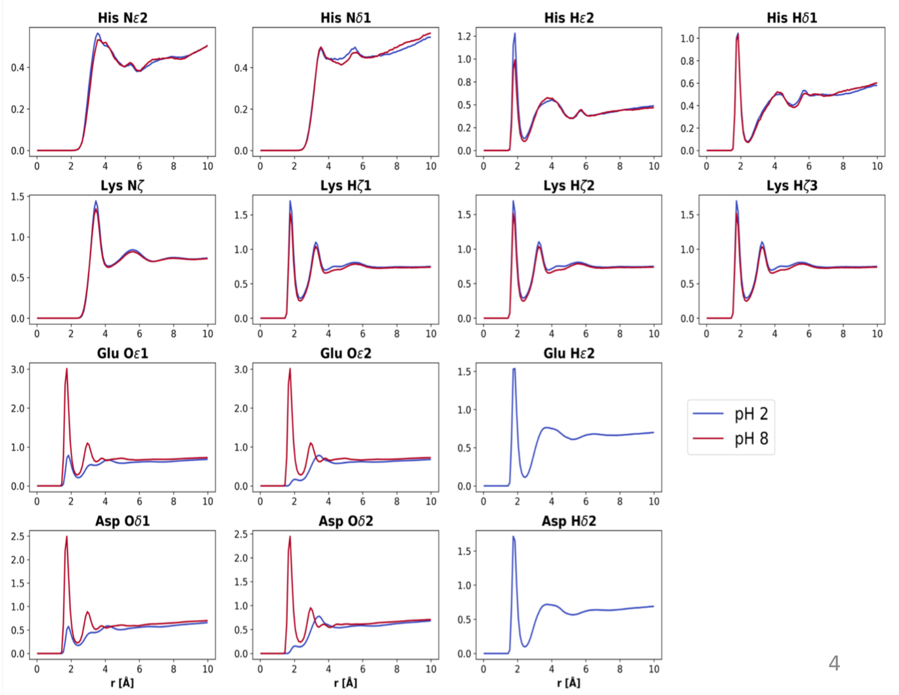

Figure S4. The distribution of inter-subunit Cα distances for Tyr419 at pH 5 (green) and pH 8 (orange) from ECpH simulation.

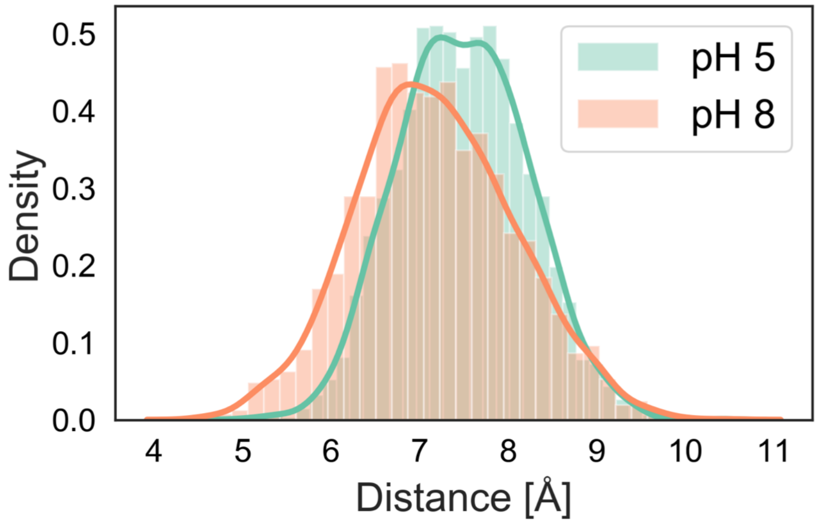

Figure S5. The distribution of RMSD of G, N, O and P helices referenced to PDB 1OTS from 0.5 $\mu$s HREMD ECpH simulation at pH 5 for two subunits of CLC-ec1 dimer. The RMSD values were calculated for Ca atoms of residues 170 – 201 (helix G), residues 355-379 (Helix N), residues 385 – 401 (Helix O) and residues 404 – 417 (Helix P).

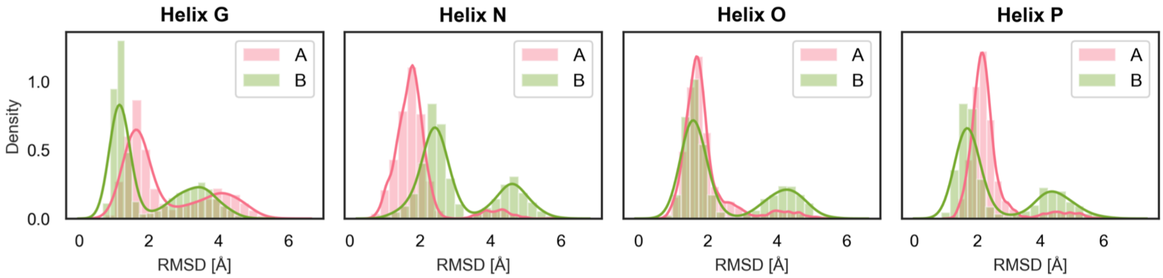
